# Genetic Gains in IRRI’s Rice Salinity Breeding and Elite Panel Development as a Future Breeding Resource

**DOI:** 10.1101/2023.06.14.544895

**Authors:** Apurva Khanna, Joie Ramos, Ma Teresa Sta. Cruz, Margaret Catolos, Mahender Anumalla, Andres Godwin, Glenn Gregorio, Rakesh Kumar Singh, Shalabh Dixit, Jauhar Ali, Md Rafiqul Islam, Vikas Kumar Singh, Akhlasur Rahman, Hasina Khatun, Daniel Joseph Pisano, Sankalp Bhosale, Waseem Hussain

**Affiliations:** Rice Breeding Innovation Platform, International Rice Research Institute (IRRI), Los Baños, Laguna 4031, Philippines; Southeast Asian Regional Center for Graduate Study and Research in Agriculture (SEARCA) and University of Philippines, Los Baños, Laguna 4031, Philippines; Crop Diversification and Genetics, International Center for Biosaline Agriculture (ICBA), 14660, Al Ruwayyah 2, Academic City, Dubai, United Arab Emirates; Plant Breeding Division, Bangladesh Rice Research Institute (BRRI), Gazipur 1701, Bangladesh; IRRI South Asia Regional Center (IRRI-SA Hub), Hyderabad, Telangana 502324, India

**Keywords:** Rice, Historical Data, Pedigrees, Salinity, Genetic trends

## Abstract

Genetic gain is a crucial parameter to check the breeding program’s success and help optimize future breeding strategies for enhanced genetic gains. In this work, IRRI’s historical data from the Philippines and Bangladesh of the salinity breeding program was used to estimate the genetic gains and identify the best lines based on higher breeding values for yield as a future genetic resource. Two-stage mixed-model approach accounting for experimental design factors and pedigrees was adopted to obtain the breeding values for yield and estimate genetic trends under the salinity conditions. A positive genetic trend of 0.1% per annum with a yield advantage of 1.52 kg/ha for the Philippines and 0.31% per annum with a yield advantage of 14.02 kg/ha for Bangladesh datasets was observed. For the released varieties, genetic gain was 0.12% per annum with a yield advantage of 2.2 kg/ha/year and 0.14% per annum with a yield advantage of 5.9 kg/ha/year, respectively. Further, based on higher breeding values for grain yield, a core set of the top 145 genotypes with higher breeding values of >2400 kg/ha in the Philippines and >3500 kg/ha in Bangladesh with a selection accuracy >0.4 were selected for formulating the elite breeding panel as a future breeding resource. Conclusively, higher genetic gains are pivotal in IRRI’s rice salinity breeding program, which requires a holistic breeding approach with a major paradigm shift in breeding strategies to enhance genetic gains.

**Key Message:** Estimating genetic gains and formulating a future salinity elite breeding panel for rice pave the way for developing better high-yielding salinity tolerant lines with enhanced genetic gains.

## Introduction

Rice (*Oryza Sativa L*.) is a major staple food crop, particularly in Asia, Latin America, and Africa. Rice is the most sensitive to soil salinity (EC above four dS/m). Millions of hectares of land in South Asia, Southeast Asia, and Africa are adopted for rice cultivation but have lower yields due to salinity stress effects (Smajgl et al. 2015). The future rice food security heavily depends on the rapid development of high-yielding salinity tolerant lines with much better adaptation to the changing climatic scenarios.

The salinity breeding program at IRRI has been at the forefront of developing salt-tolerant rice varieties utilizing various donor lines and landraces following conventional breeding approaches (Negrao et al. 2011). In the last 2-3 decades, immense efforts have been made at IRRI to develop high-yielding salt-tolerance rice varieties through conventional and molecular breeding approaches. The salinity breeding program at IRRI was further boosted by the STRASA (Stress Tolerant Rice for Africa and South Asia) Project launched in the year 2005 and continued up to 2019 (https://strasa.irri.org/varietal-releases/salinity-iron-toxicity) and Green Super Rice (GSR) projects (2006-2016). The progression of these projects led to the identification of new donor lines for vegetative and reproductive stage tolerance and the dissemination of more than 50 varieties for cultivation in saline coastal, saline, and irrigated areas/ecosystems (https://strasa.irri.org/varietal-releases/salinity-iron-toxicity, Ali et al. 2012, Singh et al. 2021). The varieties developed through these projects offer great potential for cultivating them in saline environments to increase rice production.

In the future, rice production will be immensely limited by extreme environmental conditions of salinity which would exacerbate due to climate change (Liu et al. 2020). For example, in the coastal regions of South and Southeast Asia, a source of 65% of global rice production, an increase in flood and salt intrusion has been found due to direct consequences of climate change (Radanielson et al. 2018). In 2050, the human population will reach 10 billion, and the demand for rice production will increase by 87% (Solis et al. 2020, Rawat et al. 2022). Due to the limited resources and less availability of land in the future, meeting the future rice demands will be a daunting challenge. Moreover, future rice production will only be met with heavy reliance on irrigation water (Liu et al. 2020). However, dependency on irrigation water for rice production comes with an additional cost of land salinization, and the level of dissolved salts in irrigation water has significantly increased in the past 20 years (Liu et al. 2020). Thus, global rice food security mainly depends on plant breeders to develop high-yielding, salinity-tolerant lines with broader adaptation to the above climatic changes. To develop better salinity-tolerant lines with wider adoption and achieve the required food demands in challenging conditions, it is critical to check the progress of the existing salinity breeding program and assess where we stand and how we can move forward for better improvement and enhanced genetic gains. Genetic gain is an important parameter to check the progress of the breeding program and measure its efficiency. The breeding program’s achieved rate of genetic gain will immensely help guide future breeding strategies and help allocate resources and rapid development of varieties for enhanced genetic gains. Genetic gains under salinity environments at the global level in rice have never been estimated.

Thus, this study was undertaken to accomplish two primary objectives: (i) estimating the genetic trends in the IRRI’s salinity breeding program using the data from IRRI, Philippines, and Bangladesh, and (ii) identifying top-performing genotypes based on high grain yield breeding values as future breeding resources.

## Materials and methods

### Breeding materials and experimental details

For this work, the historical datasets from salinity breeding trials conducted at various locations in Philippines from 2008 to 2019 (12 years) and Bangladesh from 2005-2014 (10 years) were used. The major traits of focus were grain yield (kg/ha) and days to flowering. The Bangladesh trials were undertaken in the districts Satkhira (22.7185° N, 89.0705° E), Ghazipur (25.5878° N, 83.5783° E), and Rajshahi (24.3745° N, 88.6042° E). The trials were organized twice a year, in two season’s dry and wet seasons in the Philippines and Aman and Boro in Bangladesh. The genotypes were staggered based on their maturity groups; early, medium, and late to synchronize appropriate stress imposition. The genotypes were screened for tolerance to salinity stress across trials starting tillering onwards or at the reproductive stage. The genotypes were planted in customized saline micro plots for imposing salinity stress, and the standard protocol was used to screen for the salinity stress. The experimental designs in the trials conducted in the Philippines varied across years from randomized complete block design (RCBD), row-column design, augmented RCBD, and alpha-lattice; however, all the trials conducted in Bangladesh were organized in RCBD.

### Pre-processing and Quality Check of the Data

The breeding values were estimated yearly, taking season and location combinations as a single trial or environment. The historical datasets retrieved were subjected to pre-processing and quality checks to ensure high-quality trials and phenotypes are retained for the downstream analysis and estimating the breeding values and genetic gains. The data pre-processing was done per the procedure detailed in Khanna et al. 2022a and Hussain et al. 2022. Trials with unexpected phenotypic values, high missing data points (>20%), missing replications, and/or design errors were filtered. After filtering, the trials were subjected to quality checks by removing the extreme data points and outliers using the Bonferroni-Holm test for studentized residuals (Bernal-Vasquez et al. 2016; Philipp et al. 2019). After pre-processing and quality check, the dataset consisted of 86 trials with 16,251 phenotypic data points with 4993 unique genotypes from IRRI HQ Philippine data sets. For Bangladesh, 110 trials possessing 3097 data points with 600 unique genotypes were retained. The details of the trials conducted across the two countries are outlined in Supplementary Table 1.

### Retrieval of pedigrees and crossing strategies

The pedigree data consisting of the parent’s and grandparent’s information on the 4,993 genotypes was utilized for substituting the pedigree-based-relationship matrix in the Philippines dataset only. Pedigree information was not available for the data sets from Bangladesh. The information of grandparents up to ten generations, along with the crossing strategy employed for each genotype, was retrieved from the state of art repository, B4R (Breeding 4 Results (B4R), 2021, https://b4r.irri.org) and IRRI genealogy management system (McLaren et al. 2005; Collard et al. 2019) with their customized R scripts. The genotypes were bred across the years employing various breeding strategies, single, double, three-way, complex crosses, and backcrosses based on their breeding objectives. The pedigree information of the genotypes for Bangladesh was unavailable and hence was not accounted for in estimating the breeding values.

### Statistical modeling

Due to different experimental designs across the trials and to account for the specific experimental design factors, the two-stage approach of mixed model analysis (Piepho et al. 2008; Piepho et al. 2012; Smith and Cullis 2018) was used. The two-stage approach also reduces the time and computational burden of analyzing huge datasets (Smith et al. 2005). In the first stage, adjusted means or best linear unbiased estimates (BLUEs) per year for each genotype were extracted for grain yield. The mixed model consists of genotypes as fixed effects with replications and seasons as random effects. Days to flowering (DTF) was used as a covariate in the model to reduce error owing due to the difference in the synchronization of flowering and to ensure the selections for tolerant genotypes would be across different maturity groups. The strategy of covariance adjustment of DTF would significantly reduce variance due to differentiation in flowering time among the genotypes in the analysis (Moreno-Amores et al. 2020; Juma et al. 2021; Khanna et al. 2021a, b). The model used in the first stage of analysis is given below:

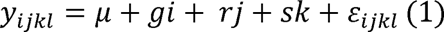

where, *y_*ijkl*_* represents BLUEs or phenotypic observation for the traits, this is with respect to each datapoint or genotype classified as the ith observation in jth replication, and kth season, *μ* is the overall mean, *g_i_* is the fixed effect of ith genotype, *r_j_* is the random effect of jth replication in each trial, *s_k_* is the random effect for season and *ɛ_*ijkl*_* is the residual error. The random effects were independently and identically distributed (IID).

In the second stage, the BLUEs estimated from the first stage were weighted and used as a response variable (Damesa et al. 2017, Khanna et al. 2022a). The weights were estimated by calculating the inverse of the squared standard errors (Mohring and Piepho 2009), which minimized the heterogeneous error variance. In this stage, a relationship matrix based on the pedigrees was fitted to account for the genetic covariances among the genotypes for reliable estimates of breeding values. The same model was used for the Bangladesh data set without fitting the pedigree matrix to extract the BLUPs. The model fitted in the second stage is as follows:

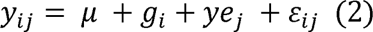

where *y_*ij*_*is the BLUE values weighted by the standard errors for *i*th observation in *j*th year, *μ* is the overall mean, g is the breeding value of *i*th genotype with g_i_ ∼ N (0, A ^2^) where ^2^ is the *i* σ g σ g genetic variance and A is the additive genetic pedigree relationship matrix, ye_*j*_ is the random effect of year, and ɛ_*ij*_ is the residual error, with ɛ_*ij*_ ∼ N(0, Rσ2ε), where R is the identity error covariance matrix and σ2ε is the error variance. The reliability values (Isik et al. 2017) of the breeding values with respect to each genotype were calculated using the equation mentioned below:

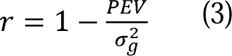

Heritability for the yield was estimated using the method suggested by Cullis et al. 2006 and Piepho and Mohring 2007 using model 1 with genotypes as a random effect. This approach is useful when the data is highly unbalanced, with uncommon genotypes screened across the years and seasons. The following equation was used to calculate the heritability for trials per year:

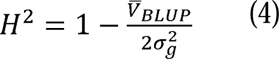

All the analysis was done using the ASReml-R package (Butler et al. 2017) in the R software (R Core Team, 2020). The pedigree-based relationship matrix (A-matrix) was constructed using the R package AGHMatrix (Amadeu et al. 2016).

### Estimation of the genetic trends

For the IRRI-HQ data, the genetic gains were estimated by regressing each genotype’s breeding values over the year of origin or the year when the cross was attempted for each genotype. The year of origin for each genotype record was extracted using the customized R scripts from the genealogy management system IRRI (McLaren et al. 2005; Collard et al. 2019). However, in the case of Bangladesh, the genetic trends were estimated by regressing the BLUPs over the year of testing for each genotype. To estimate the gains only with released varieties, a similar strategy was followed by regressing each genotype’s breeding values or BLUPs over the year of release for each of the two countries. Additionally, genetic gain trends were plotted using the non-linear approach of *loess* (Local weighted regression) to check the short-term and long-term genetic trends in the salinity breeding program at IRRI, Philippines, and BRRI, Bangladesh data sets.

### Formulation of elite breeding panel

Breeding values or BLUPs for grain yield obtained from the second-stage analysis were used to formulate the salinity breeding panel as a future genetic resource. Based on the higher breeding values of >2300 kg/h in the Philippines and BLUPs >3550 kg/h in Bangladesh and prediction accuracy of >0.4, 145 genotypes were selected as a part of the breeding panel. For the Philippine IRRI-HQ data, genetic similarity between the selected lines and in comparison, to the whole historical line collection was assessed using the relationship matrix based on pedigrees. The diversity and similarity of the lines over the complete set of 4993 genotypes from the historical salinity dataset was visualized through the bi-plot graph. The variables for the bi-plot were obtained through the principal component analysis (PCA) performed using the function *princomp* in R software on the A-matrix or pedigree matrix.

### Genetic Trends of released lines

From the IRRI-HQ data in the Philippines, 17 IRRI-released saline tolerant varieties were utilized to estimate the genetic gains. Similarly, the gains were estimated using 12 released salinity-tolerant varieties in Bangladesh. The breeding values from the Philippine data set and BLUPs from Bangladesh data were regressed to their year of release for estimating the genetic gains. Also, to further understand the breeding program’s growth, recently nominated 25 IRRI varieties across nine countries for the year 2021-22 were compared for their breeding values.

Superior performing 12 genotypes comprising seven nominated varieties and four selected varieties from the historical core panel possessing higher breeding values and salinity tolerance were tested for stability using their grain yield performances in 5 environments, *viz.,* in the year 2018 at IRRI, Philippines, and Ajuy Iloilo during the wet season: in the year 2019 at IRRI Philippines during dry and wet seasons and at Ajuy Iloilo during the wet season. Stability analysis was performed using R software’s GGE Biplot GUI package (Frutos et al. 2014). The percentages of GGE explained by the top two PC axes were estimated for ranking genotypes based on their relative performance and ranking genotypes relative to the ideal genotype. The analysis was based on a Tester-centered (G+GE) table without scaling and with row metric preserving.

## Results

### Description of historical salinity datasets

A high difference in the mean values for grain yield (kg/ha) was observed in the Philippine and Bangladesh data sets (Fig. 1 a, b). For the DTF in the Philippines dataset, two maturity categories of early and medium were found, with 98% (DTF: 66-109 days) of genotypes falling under early and 2% (DTF: 110-124 days) genotypes under medium maturity groups. However, in the Bangladesh dataset, the genotypes were found to fit into all three maturity groups, with 66% of genotypes possessing DTF values between 70-109 days, 27% of genotypes having DTF ranges between 110-124 days, and 7% of the genotypes were the late category with DTF ranges between 124-135 days respectively.

**Fig. 1.**
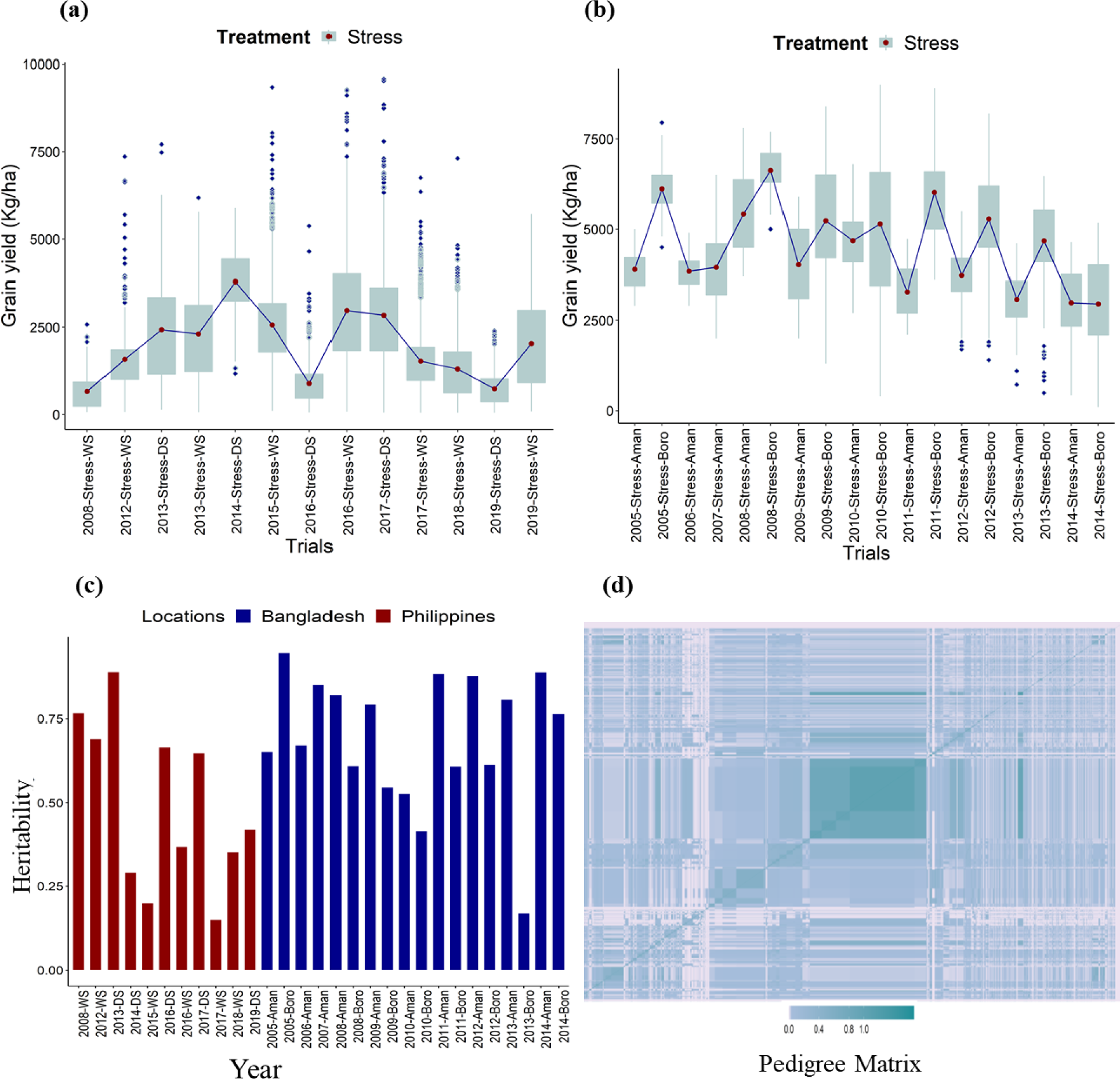
Boxplots depicting the ranges of raw grain yields (kg/ha) across historical breeding trials for salinity stress from **(a)** 2008-2019 at the Philippines, **(b)** 2005-2014 in Bangladesh. The grain yields have been shown season-wise **(a)** DS and WS, symbolize dry and wet seasons for the mentioned respective years at the Philippines; likewise in **(b)** aman and boro, represent the trial seasons in Bangladesh respective to mentioned years. **(c)** The bar plot shows the heritability of the trials at IRRI, Philippines, from 2008 to 2019, as depicted by the red bars, and the heritability of the trials conducted in Bangladesh between 2005 to 2014, as depicted by the blue bars. The WS and DS represent the wet and dry seasons in the Philippines. Similarly, aman and boro are two seasons in which trials were conducte in Bangladesh **(d)** Pedigree matrix depicting pedigree-based relationship based on 4,993 unique genotypes bre across 12 years at IRRI, Philippines. The range is represented by blue and green color, As the color scale progresses from light blue to through sea green in the off diagonals to dark cyan in the diagonals, the genetic similarit increases amongst the genotypes scaling 0.4 to 0.8 to 1. The higher the genetic similarity the higher the score.

The heritability based on BLUP differences for grain yield was estimated season-wise for each year. The heritability estimates ranged between 0.20-0.89 for the trials from 2008-2019 in the Philippines dataset (Fig. 1c). Similarly, in the historical dataset from Bangladesh, the heritability ranges were between 0.17-0.95 (Fig. 1c). The heritability values were very low in few of the seasons due to the environmental/season/year influence of genotypes to the salinity stress conditions, which would, in turn, affect the grain yields (Rauf et al. 2012). This could be due to different stress levels and environmental conditions across the years, as the priority objective of the stress trials would be to impart higher salinity stress for identifying tolerant genotypes.

### Historical data connectivity

The connectivity in the historical dataset is a major parameter affecting the estimates of genetic gains. The current study had apt connectivity in the data across growing seasons and years. In the Philippine data, the pedigree matrix was included in the second stage of analysis to build the connectivity (Khanna et al. 2022a) and get a reliable estimation of the breeding values and hence genetic gains (Fig. 1d). Additionally, the suitable connectivity in the data can be attributed to the common saline tolerant (FL478, Pokkali-8558) and susceptible checks and varieties (IR29, IRRI 104, IRRI 165, etc.,) used in the breeding trials across seasons and years. Alongside, the connectivity was maintained as the tolerant genotypes/varieties were evaluated in subsequent years to reconfirm their stable tolerance across years (Fig. 2a, b). It was also observed that there was connectivity amongst the trials across seasons in each year in both countries (Supplementary Fig 1a,b,c,d)

**Fig. 2.**
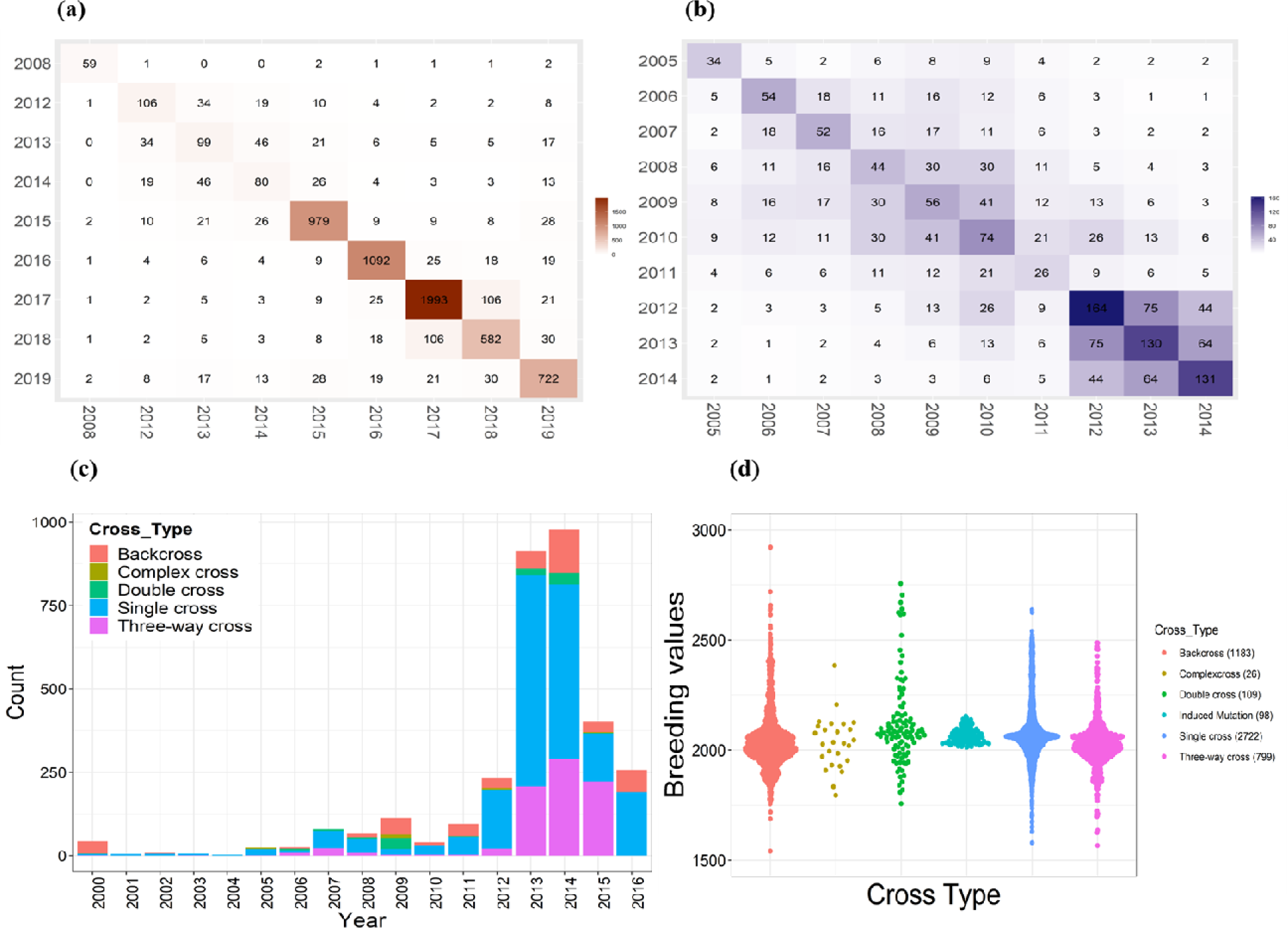
**(a)** Connectivity of all the unique genotypes in the Philippines dataset, **(b)** Bangladesh dataset across years were plotted in form of heatmap. The numbers in each box represent common genotypes between each of the year combinations. Since the checks and promising varieties were repeatedly bred and tested in the successive years for evaluating their performance, the datasets represent apt connectivity across years. **(c)** The barplot depicts the classification of the genotypes tested across years of origin in the trials organized at IRRI, Philippines from 2000-2016. The differential coloration in each bar represents varying breeding strategies employed in the respective years. As the figure depicts single and backcrosses were initiated in the initial years which were added by double three-way complex crosses in the later years in the breeding program. **(d)** The boxplot depicts the range of each of the breeding strategies in terms of breeding values. However, the ranges slightly differ in each of the cases, the backcrosses possessed a wider range followed by single and three way crosses. The numbers of each of the cross combinations formulated across years have also been depicted in the brackets besides each cross type. The maximum were single cross combinations followed by backcrosses.

### Crossing strategy across salinity historical breeding program

It is crucial to decipher the crossing strategy or the breeding scheme adopted by the breeders during this period and associate it with the genetic gain trends. The crossing strategy used was extracted from the B4R database. From 2000 to 2016, the breeders performed single, double, three-way, complex, and backcrosses (Fig. 2 c, d). It was found that during the initial years, 2000 to 2005, most of the crosses were single and backcrosses, most of which, until 2012, included double, three-way, and complex cross combinations. However, post-2012, the era when IRRI was rephrasing from marker-assisted backcross breeding to complex/multiple crossing strategies for developing various stress-tolerant breeding varieties, the complex crossing strategy was more highlighted along with three-way and backcrosses (Fig. 2c). The unique 4, 993 genotypes from the historical breeding trials dataset of Philippines had a broad background with differing parents and cross combinations and can also be classified into 770 families based on their diverse parental crosses, amongst which 108 families were derived from backcrosses; 451 from single cross combinations; 49 from double cross combinations; 152 from three-way crosses; 13 from complex crosses, two from induced mutations procedures and remaining were landraces, and remaining 5 were accessions and donors from the gene bank. However, no clear association was found between the crossing strategy and change in the genetic gain trends. After 2012, more fluctuations were observed with genetic trends because the breeding program focused on complex/multiple crossing strategies to develop multiple stress-tolerant breeding lines with a limited focus on using population improvement strategy to enhance the yield.

### Estimation of breeding values and genetic trends

The breeding values ranged between 1326.06-4720.35 kg/ha in the Philippines dataset and 3587.27-5829.77 kg/ha in the Bangladesh dataset (Supplementary Fig. 2a, b). The genetic gain at IRRI, Philippines, between 2008 and 2019 was 0.1% per annum, with a yield advantage of 1.53 kg/ha/year. The gain was estimated by regressing the year of origin or year of crossing for each of the genotypes spanning from 2000 to 2016, tested in the breeding trials across the years 2008 to 2019 (Fig. 3a). In Bangladesh, the genetic gain was 0.31% per annum with a yield advantage of 14.02 kg/ha per annum (Fig. 3b). Both in IRRI Philippines and BRRI, Bangladesh data sets, linear and non-linear genetic trends were plotted to check the fluctuations in genetic trends over the years (Fig. 3a, b). The genetic trends were estimated for the released varieties in the Philippines and Bangladesh. The genetic gain was 0.12% per annum with a yield advantage of 2.2 kg/ha/year as estimated for the released varieties at IRRI, Philippines (Fig. 3c). The gain was 0.14% per annum with a yield advantage of 5.9 kg/ha/year (Fig. 3d).

**Fig. 3.**
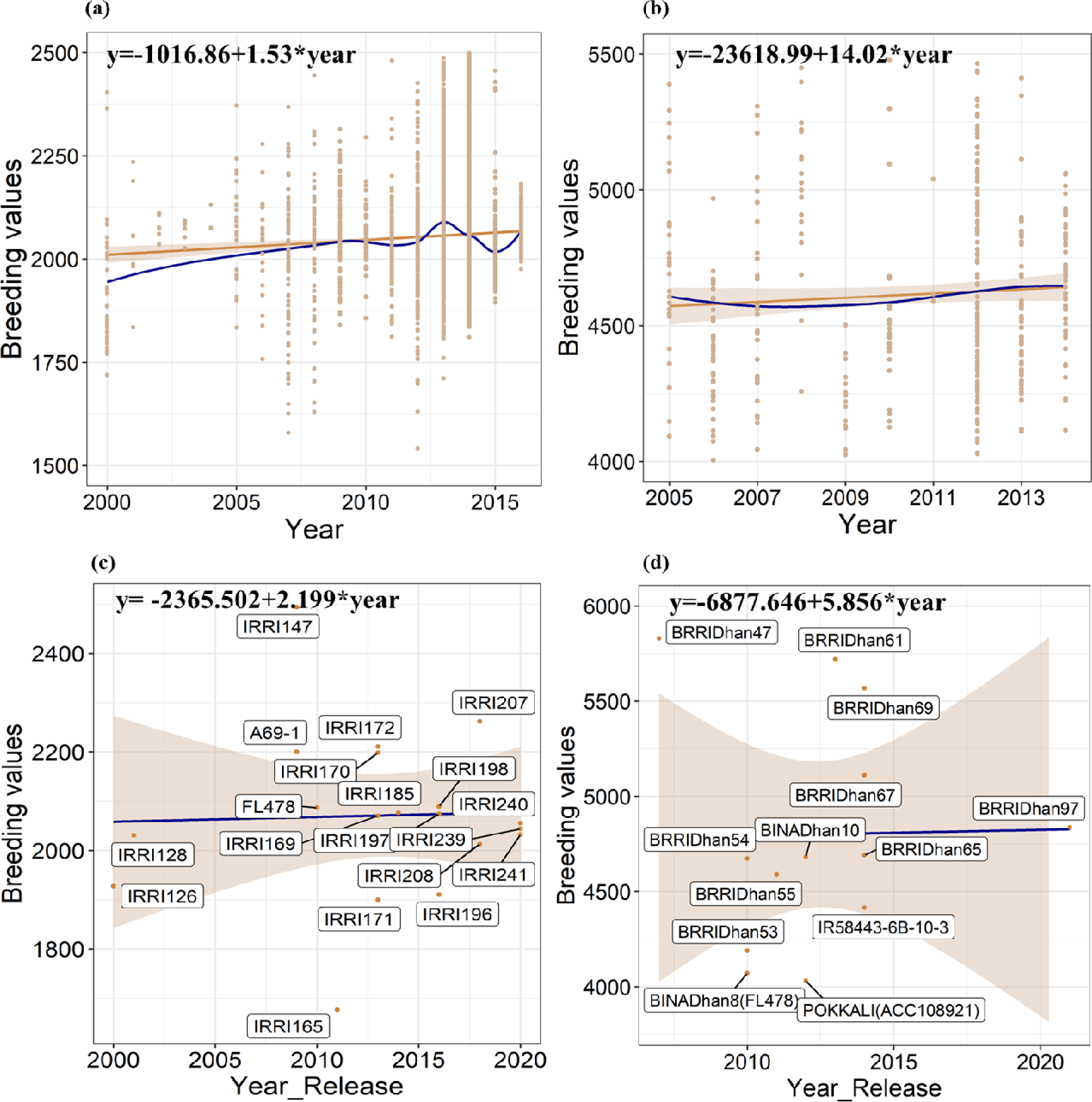
**(a)** Trends in genetic gain from IRRI’s salinity breeding program comprising 12 years of breeding trials, 2008 to 2019. **(b)** Trends in genetic gain from salinity breeding program in Bangladesh comprising 10 years of breedin trials, 2005 to 2014. In both (a) and (b), The x-axis depicts the year of origin, and the y-axis portrays the breedin values or BLUPs of the genotypes. The dots represent the breeding values or BLUPs respective to each year. The slope in peru represents the genetic trends using linear regression, and the slope in dark blue portrays the genetic trend using a non-parametric approach using loess regression. **(c)** The trends in genetic gain for the released varieties across years bred for salinity tolerance at IRRI. **(d)** The trends in genetic gains for the released varieties across years bred in Bangladesh. The gains were likewise estimated by regressing the grain yield breeding values on the year of origin.

### Development of an elite breeding panel

The genotypes with higher breeding values of >2400 kg/ha in the Philippines and >3500 kg/ha in Bangladesh and having selection accuracy >0.4 were selected for formulating the elite breeding panel. The top 145 genotypes were selected as a future breeding resource for the elite core panel. The criteria for selecting the elite lines, their breeding values, cross combinations, and crossing strategies employed are given in Supplementary Table 2. We also accessed the kinship of the lines from IRRI-HQ Philippine data using the pedigree relationship matrix (Fig. 4a). The genotypes selected for the elite core panel represent the whole data collection and cover the diversity of the entire collection very well. The genotypes selected for the elite panel not only possess a high breeding value for yield but harbor salinity-tolerant landraces, including Sadri, Pokkali, and Cheriviruppu; elite breeding genotypes like NERICA (New Rice for Africa), which are early maturing (<100 days); tolerant to major stresses of Africa; AT401, variety can withstand coastal saline environments; Zinc fortified genotypes (IR68144; BR7840-54-3-1); Zinc efficient donor parents (IR55179). The panel additionally possesses genotypes with superior characters, including genotypes with superior yields under DSR conditions (Supplementary Table 3), zinc-efficient genotypes with superior breeding values, iron toxicity tolerant genotypes, and coastal and acid saline tolerant genotypes (Supplementary Table 2).

**Fig. 4.**
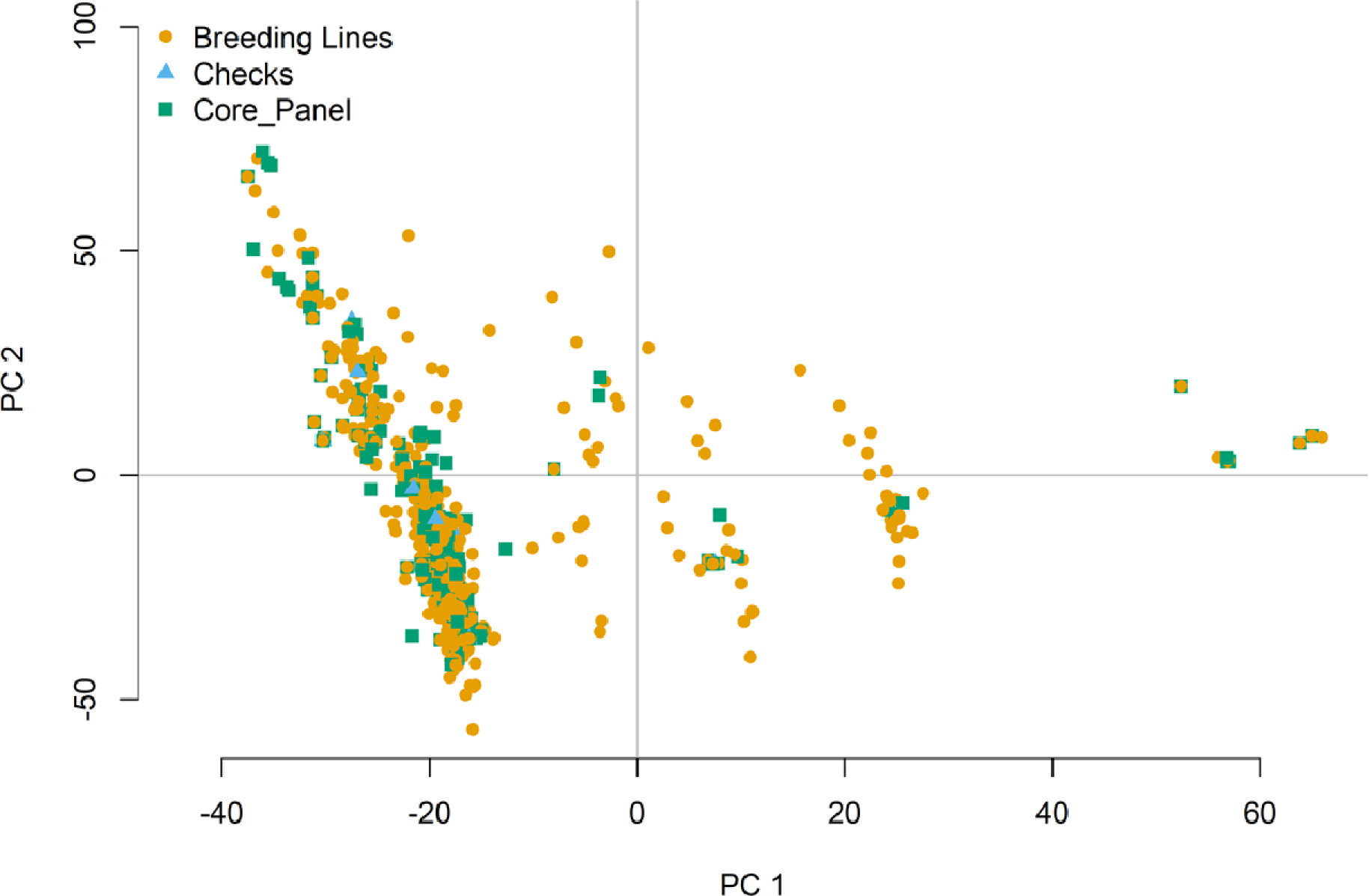
PCA-Biplot depicting the complete set of unique breeding lines obtained from the historical breeding trials in IRRI-HQ. The elite core panel breeding lines selected based on the superior breeding values have been depicted by violet color and checks by blue color. The biplot was obtained and plotted using the first two principal components using the pedigree matrix. The elite core panel lines overlap the complete set of breeding lines and represent the diversit of whole collection as a useful future breeding resource.

### Comparison of breeding values in IRRI released, nominated, and core panel lines

The breeding values for the IRRI-released varieties were compared with the saline tolerant checks and varieties for estimating the gains obtained in the released varieties. The comparative analysis would help to comprehend the genetic progress and identify superior genotypes across the years. The breeding values of the released varieties were superior to the popular checks for both countries. In the Philippines dataset, the released global salinity varieties IRRI 147, IRRI 207 and IRRI 172, IRRI 170, and IRRI 198 depicted superior breeding values of 2493.84 kg/ha, 2261.63 kg/ha, 2210.60 kg/ha, 2198.40 kg/ha and 2089.28 compared to other salinity tolerant varieties and popular checks A69-1 and FL478 with the breeding value of 2200 kg/ha and 2086.74 kg/ha respectively. Amongst these superior-performing varieties, IRRI 207 and IRRI 198 were recently released in the years 2018 and 2016 (Fig. 2c). In the Bangladesh dataset, all the genotypes included for estimating the gain were superior performing to salinity tolerant check FL478. Amongst all, BRRI Dhan 47, BRRI Dhan 61, BRRI Dhan 69, and BRRI Dhan 67 depicted superior breeding values of 5829.77 kg/ha, 5720.48 kg/ha, 5568.19 kg/ha and 5110.41 kg/ha respectively, to the salinity tolerant check FL478 (BINA Dhan 8) and Pokkali with the breeding values of 4071.16 kg/ha and 4021.65 kg/ha respectively (Fig. 2d). Amongst these varieties BRRI Dhan 61, BRRI Dhan 69 and BRRI Dhan 67 were released in the years 2013 and 2014 respectively.

Alongside the released varieties across years, 25 IRRI-bred varieties nominated for eight countries in 2021-22 were also compared for their breeding values. The breeding values of these varieties ranged between 2025.80 to 2920.67 kg/h (Supplementary Fig. 3). Overall, IR121094-B-B-AJY3-2-B (IR18T1021) nominated for release in Thailand depicted the highest breeding value of 2920.67 kg/h followed by IR121188-28-1-CMU2-2-B (IR18T1015), nominated for release in the Philippines with a breeding value of 2718.15 kg/h. Interestingly, the latter possesses higher breeding value compared with the till date released varieties utilized for estimating gains for 2008-2019 and amongst other nominated varieties for the year 2021-22 for the Philippines. Additionally, IR121188-28-1-CMU2-2-B was also found to be the most stable variety based on the relative performance when ranked amongst the other varieties, including IRRI 147, which possessed the highest breeding value amongst the date-released varieties for the Philippines, and was found the most preferred variety residing along relative to the ideal genotype as shown by the arrow in the Supplementary Fig. 4.

Further, succeeding IR117676-318-1-1-1 has been nominated for Sri Lanka, which possesses a breeding value of 2640.39 kg/h and ranked fourth for stability amongst the tested varieties, followed by IR112462-B-25-2-1-1 (IR16T1631) nominated for Bangladesh and Lao PDR with a breeding value of 2625.29 kg/h. IR16T1009, nominated for Thailand, also depicted a breeding value of 2514.73 kg/h. This was followed by IR63307-4B-4-3, nominated for Indonesia, Vietnam, and Sri Lanka with a breeding value of 2493.843 kg/h, bred using a soma clonal variant of Pokkali. Another variety, IR117839-22-15-B-CMU10-1-B, nominated for release in the Philippines and Vietnam, possessed a breeding value of 2479.29 kg/h and was the second most stable genotype amongst the tested varieties in the five environments.

In the selected panel, 8% of the genotypes comprise IRRI-bred varieties that can withstand salinity stress in hand with superior performance under DSR conditions with yields ranging between 2195-4758 kg/h (Supplementary Table 3). Regarding including the genotypes from current year nominations, 12% of the panel comprises recently nominated varieties with the highest breeding values. In all eight superior breeding value harboring genotypes of the nominations *viz*., IR121094-B-B-AJY3-2-B (2920.67), IR121188-28-1-CMU2-2-B (2718.15), IR117676-318-1-1-1 (2640.39), IR112462-B-25-2-1-1 (2625.29), IR16T1009 (2514.73), IR63307-4B-4-3 (2493.843), IR117839-22-15-B-CMU10-1-B (2479.29), IR117749-B-B-CMU6-1-B (2379.65) was part of the core breeding panel (Supplementary Table 2). Another 3% of the panel formulates zinc & iron bio-fortified genotypes with superior breeding values of >2400 kg/ha for being future-ready for upscaling salinity tolerance and additional characteristics.

## Discussion

We demonstrated the genetic trends in IRRI’s rice salinity breeding program by leveraging historical data and pedigree information. Besides genetic gain estimates, top-performing genotypes based on high breeding values for grain yield were also identified as a future elite breeding resource. Availability of the pedigree information from the IRRI-HQ data was crucial to fit and use in the second stage of analysis for reliable estimation of the breeding values to help in the identification of accurate genotypes for the development of the elite panel and accurate estimation of genetic gains (Rutkoski et al. 2019a). As these lines have already been bred under saline environments, they are not only tolerant to salinity but possess high breeding values for yield, making them readily available as a future genetic resource for the population improvement-based breeding strategy, further enhancing the genetic gains in the salinity breeding program.

### Genetic gain estimations

One of the main goals of the breeding program is to achieve higher and constant rates of increase in genetic gain while maintaining genetic diversity. Genetic gain estimations are crucial to assess the success and growth of the breeding programs through an increase in the mean population breeding values over the years of selection (Ramstein et.al. 2019, Rutkoski et al., 2019b). This work is the first report demonstrating the genetic gains in the rice salinity breeding program and assessing its progress in terms of genetic gains. Positive genetic gains were obtained in IRRI-Philippines and BRRI, Bangladesh. However, the rate of genetic gain was just 0.33% in Bangladesh, and 0.13% in the Philippines, which is much lower than the required rate of genetic gain in rice is 1.5% or above (Khanna et al. 2022a, Li et al. 2018). To achieve the necessary rates of genetic gains in the IRRI’s rice salinity breeding program, a major tweaking in the breeder’s equation (Cobb et al., 2019, Merrick et al. 2022) through modernization and optimization is highly required. Further, major paradigm shifts are required in rice salinity breeding to deliver high salinity tolerant lines with higher genetic gains. A highly focused population improvement with systematic pre-breeding efforts is needed to deliver constant and higher rates of genetic gains in IRRI’s salinity breeding program.

### Core panel formulation for identifying elite genotypes

A set of high-performance, elite breeding lines with salinity tolerance is highly required to unlock the potential of cultivation in saline soils with enhanced genetic gains. The conventional salinity breeding at IRRI has mainly focused on crossing the non-elite (salinity tolerant traditional donors/ landraces) to the high-yielding elite breeding lines to develop the high-yielding elite salinity tolerant lines. Further, since the inception of molecular breeding, the main focus has been on the introgression or pyramiding of salinity-tolerant QTLs in elite backgrounds (Singh et al. 2021). The rice breeders have used different crossing strategies (Figure 2 a, b, c) single, complex, double, and backcrosses to integrate these QTLs into the elite genetic backgrounds and develop the new breeding lines. Diverse materials, including landraces and donors, have been extensively used to diversify the gene pool and develop climate-resilient salinity varieties (Singh et al. 2021, Sandhu et al. 2021, Yadav et al. 2021). The genotypes used in this study represent the breadth of the diversity of IRRI’s salinity breeding program through which several high-yielding salinity tolerant varieties have been released (Singh et al. 2021). This breeding resource represents an essential source of elite genetic variation that can be leveraged to extract the diverse genotypes with high breeding values for grain yield and salinity tolerance as a future genetic resource. To this end, an effort was made to develop the representative set of the elite pool from this historical collection based on high breeding value and genetic divergence. The developed elite pool can be readily used in a rapid, recurrent selection-based breeding strategy to quickly re-cycle the lines for enhanced genetic gains. Recurrent selection is critical to increase the frequency of favorable additive alleles of grain yield and enhance genetic gains (Khanna et al. 2022a, Juma et al. 2021).

For example, the elite pool identified in this work has released lines like BRRI Dhan 55, BRRI Dhan 47, BRRI Dhan 67, IRRI 185, IRRI 235, and IRRI 147, which showed high breeding values for the grain yield (Fig. 3; Supplementary Table 2). Specifically, the released variety IRRI 147 for the Philippines, also released as BRRI Dhan 47 in Bangladesh, depicted the highest breeding values amongst the released varieties. The variety harbors a unique characteristic of erect plant architecture as its leaf angle falls between 5°-20° (BRRI annual report 2018-2019). The erect plant architecture renders higher photosynthetic abilities by impacting the source and sink organs, making it crucial to identify genotypes with superior “ideotypes,” which can significantly enhance yield, productivity, and gains (Chang et al. 2020). Additionally, genotypes IR58443-6B-10-3, IR16T1110, IR16T1086, IR16T1661, and IR16T1018 were also included in the breeding pool, and these genotypes have shown to have high performance under the direct-seeded conditions (DSR) along with salinity tolerance (IRRI personal communication). The elite pool consists of 8 genotypes from the freshly nominated 25 varieties for eight countries in 2021-22. These genotypes revealed high salinity tolerance and superior grain yield, which provide ample evidence that conscious efforts are being made to develop the high-yielding lines under saline environments to achieve desired genetic gains.

Interestingly, the nominated variety IR63307-4B-4-3 for three countries, Indonesia, Vietnam, and Sri Lanka, has been bred by crossing IR 51511-B-B-34-B/TCCP 266-2-49-B-B-3 using a single cross-breeding strategy. The donor parent used here, TCCP 266-2-49-B-B-3, is a soma clonal variant of Pokkali with superior characteristics, including semi-dwarf plant type with white pericarp, medium consistency of grain type and possesses high yield potential along with vigorous growth without lodging. All eight superior breeding values harboring nominations genotypes *viz*., IR121094-B-B-AJY3-2-B, IR121188-28-1-CMU2-2-B, IR117676-318-1-1-1, IR112462-B-25-2-1-1, IR16T1009, IR63307-4B-4-3, IR117839-22-15-B-CMU10-1-B, and IR117749-B-B-CMU6-1-B is part of the elite panel. Conclusively, the selected lines for the elite pool development don’t have only high breeding values for yield but possess a tolerance under salinity stress. The genotypes will be a great source of readily available variation to use in recurrent selection and further recombine and reshuffle in it to create an additional novel source of variation for grain yield and enhance the genetic gains. Consequently, a systematically designed holistic breeding approach was implemented with multi-location trial evaluations of the newly identified panel with significant emphasis on economically targeted traits considering the target population of the environment (TPE), plant architecture, and yield component traits (Yadav et al. 2021, Chang et al. 2020) laterally with prediction based breeding strategies would help in sustainably improving and enhancing genetic gains in the future rice salinity breeding program at IRRI (Reynolds et al. 2011; Yadav et al. 2021, Gerard et al. 2020).

## Conclusions

The current rate of genetic gains observed in the salinity breeding program is comparatively lower than the required rates of 1.5% or above. The rate of genetic gain in rice will increase to 2.5% or above in 2050. To deliver higher rates of genetic gains in the salinity breeding of IRRI, a holistic and systematic breeding effort with the integration of modern tools and technologies is required. A population improvement breeding strategy based on an elite x elite scheme with the integration of genomic selection is highly required. The elite breeding pool identified in this study would be the most potent and readily available genetic resource to drive the population improvement-based breeding strategy in IRRI’s salinity breeding program.

## Acknowledgments

We thank the Bill and Melinda Gates Foundation for funding and supporting this study. We would also like to acknowledge our respect and thankfulness to Dr. Hans Bhardwaj (Platform Leader, Rice Breeding Innovations, IRRI) for his support and encouragement. The authors also thank IRRI rainfed breeding program breeders and technical staff, who have conducted these trials since 2003 and generated the data used in this study. We would like to thank all IRRI members who have contributed to maintaining the B4R database to gather and preserve valuable breeding trial information.

## Ethics declarations

**NA**

## Additional information

The phenotypic data from the historical trials and scripts for analysis are available at the GitHub repository and can be accessed through the following links:

https://github.com/whussain2/Genetic_Trend_Rice_Drought https://github.com/whussain2/Analysis-pipeline

## Publisher’s notes

## Supplementary Information

## Supplementary Figures

**Supplementary Fig. 1.**
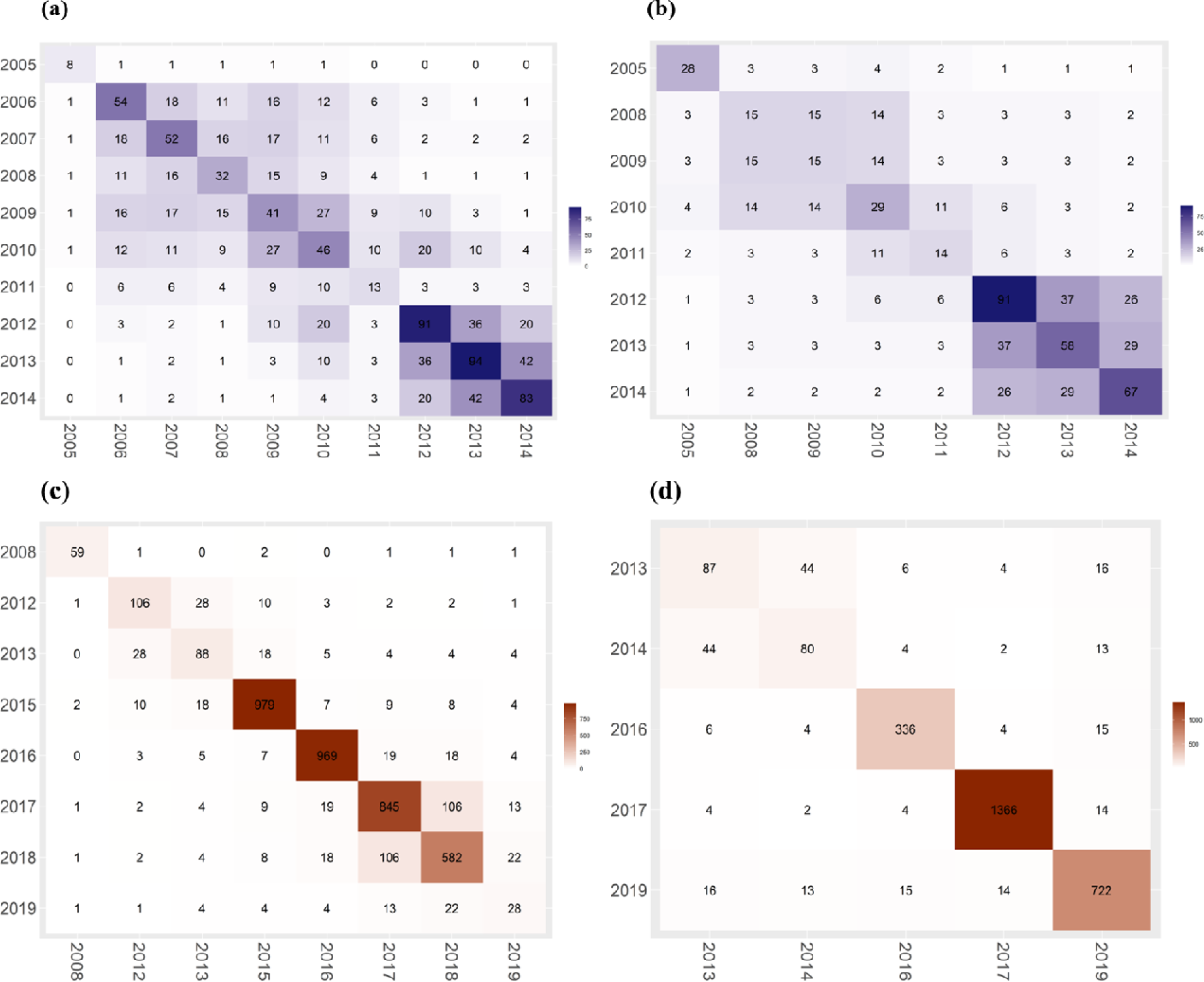
The figures depicted in brown depict the connectivity of the unique genotypes from Bangladesh evaluated during **(a)** Aman season, **(b)** Boro season; The heat maps plotted in blue depict the connectivity of the unique genotypes from the Philippines dataset tested during the **(c)** wet season, **(d)** dry season. The numbers in each box represent commo genotypes between each year combination. Both datasets represent apt connectivity across years since the checks and promising varieties were repeatedly bred and tested in successive years to evaluate their performance.

**Supplementary Fig. 2.**
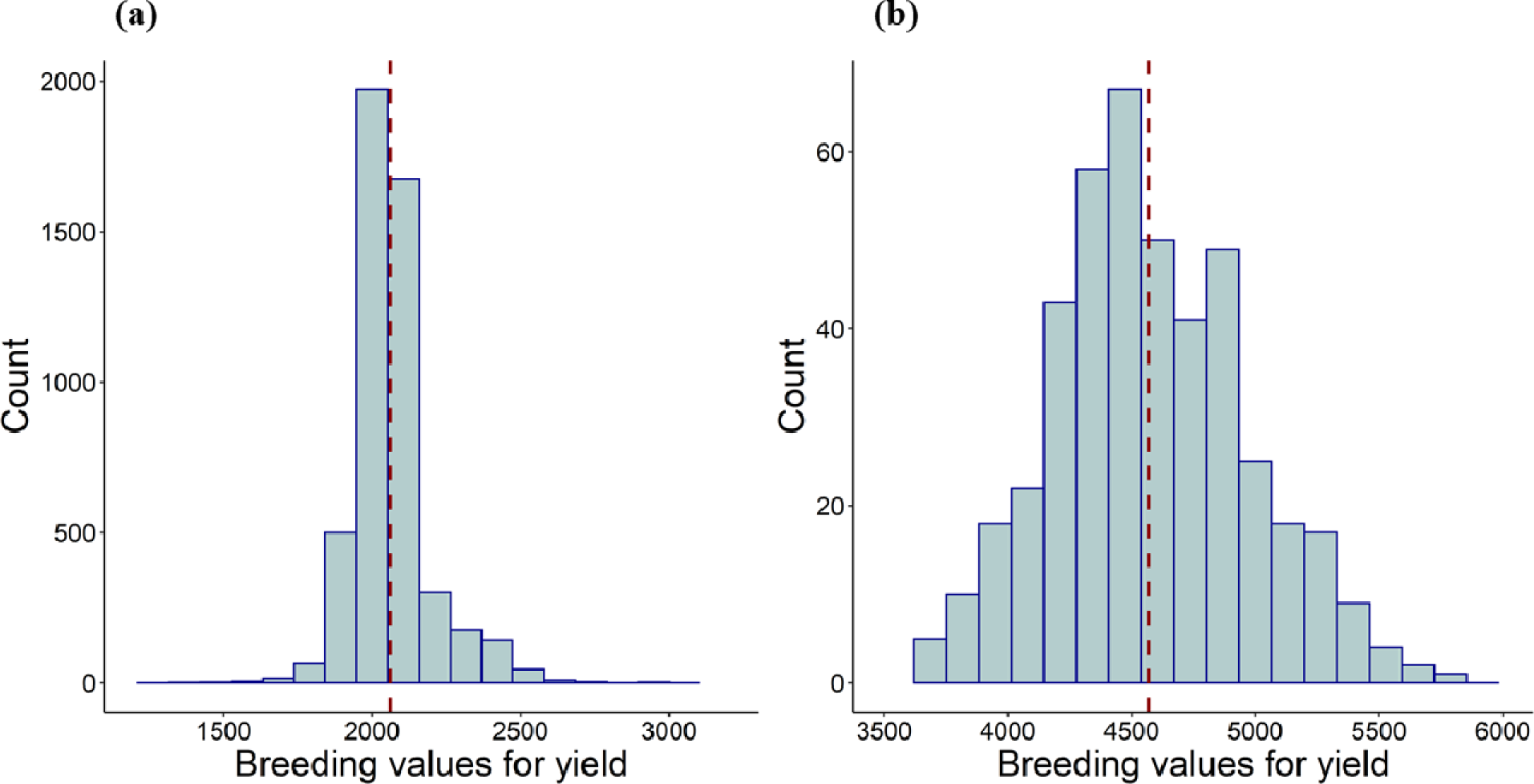
The distribution of the breeding values for grain yield (kg/ha) of the genotypes screened under reproductive stag salinity stress at **(a)** Philippines and **(b)** Bangladesh has been demonstrated. In each of the figures the red margin potrays the mean of the breeding values in each of the cases.

**Supplementary Fig. 3.**
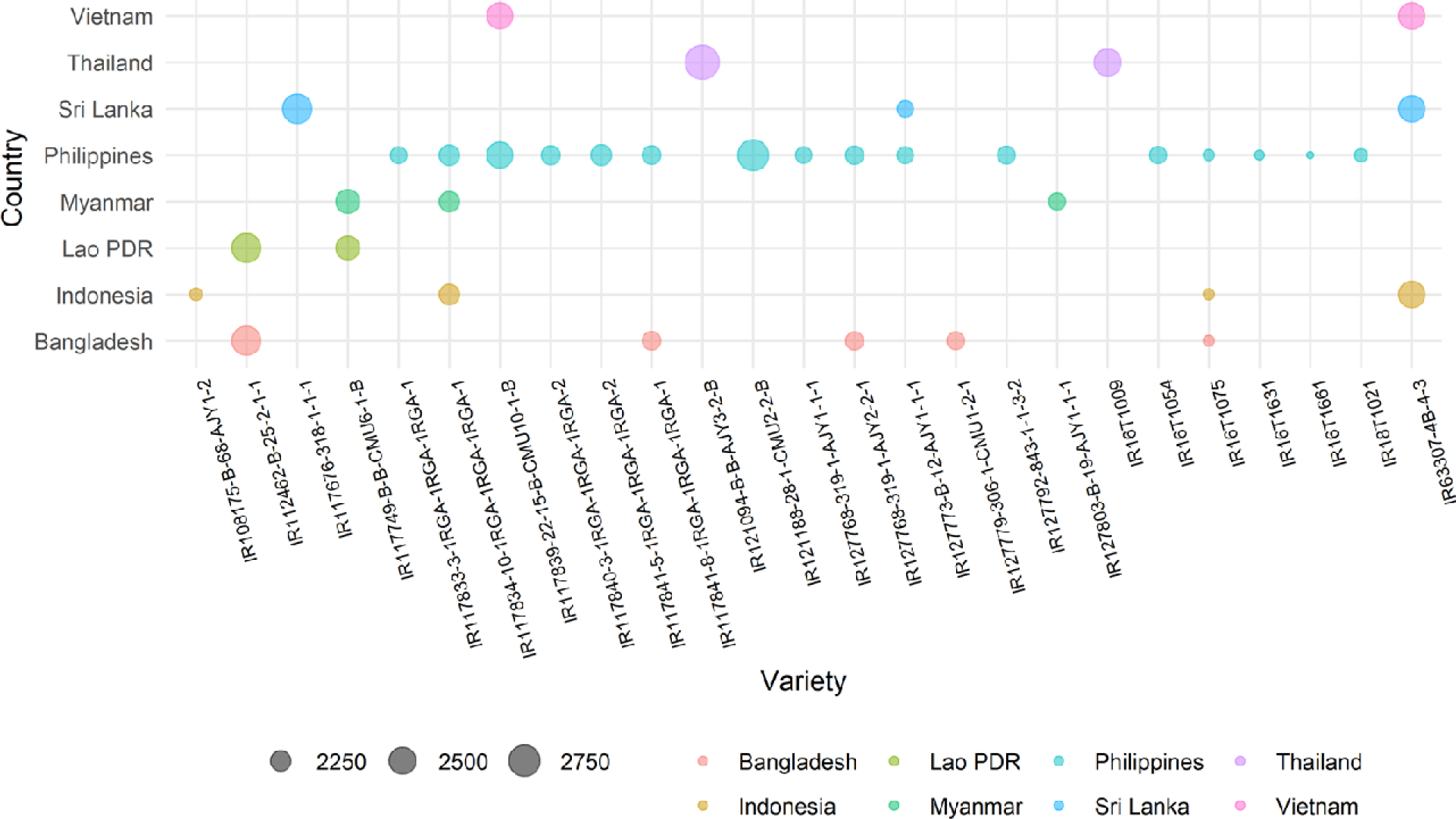
Salt tolerant historical breeding lines bred across 12 years from 2008-2019 at IRRI, Philippines and were selecte for nominations across 8 countries for the year 2021-22. The breeding values for each of the countries have bee depicted by different colors. The more the circumference of the circle, higher the breeding values. The highest breeding value was for the genotype IR121094-B-B-AJY3-2-B, nominated for Thailand depicted in violet color, followed by IR121188-28-1-CMU2-2-B, nominated for the Philippines depicted in cyan color.

**Supplementary Fig. 4.**
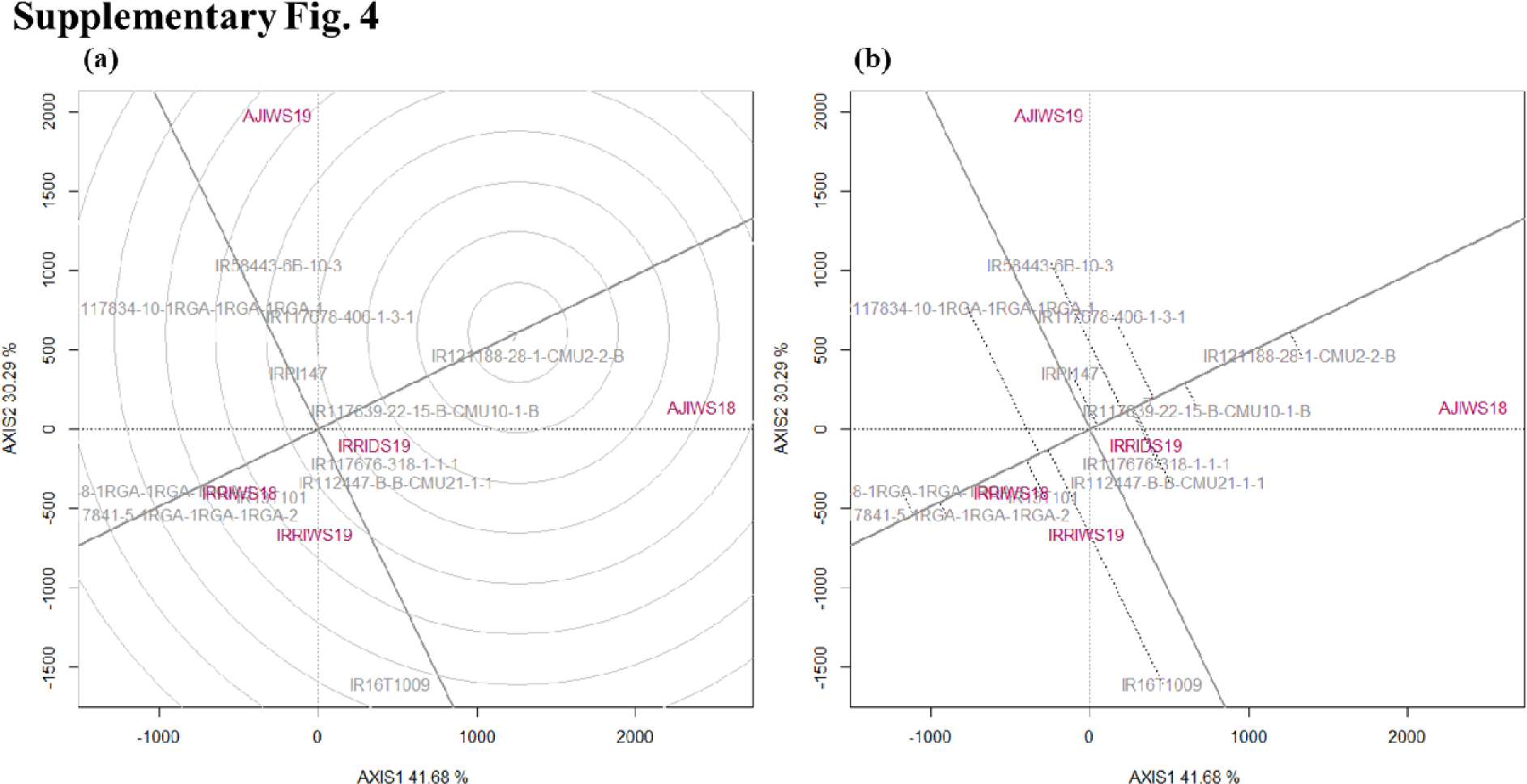
The goodness of fit of the biplot, explaining a total of 71.97% of test centric data (G+GE) has been represented in the figures. The percentages of GGE explained by the top two PC axes were estimated for ranking genotypes base on their relative performance and ranking genotypes relative to the ideal genotype. **(a)** Ranking of genotypes with reference to the “ideal genotype”, as shown by the arrow. The genotypes are raked in descending order starting from the arrow of ideal genotype. **(b)** Comparative depiction of genotypes based on their mean performance and stability portraying the average-environment coordination view of GGE biplot.

## References

Ali J, Xu J-L, Gao Y-M, et al (2017) Harnessing the hidden genetic diversity for improving multiple abiotic stress tolerance in rice (Oryza sativa L.). PLoS One 12:e0172515. https://doi.org/10.1371/journal.pone.0172515

Amadeu RR, Cellon C, Olmstead JW, Gracia AA, Resende MF, Muñoz PR (2016) AGHmatrix: R package to construct relationship matrices for autotetraploid and diploid species: A blueberry example. Plant Genome 9:3. https://doi.org/10.3835/plantgenome2016.01.0009

Bernal-Vasquez AM, Utz H-F, Piepho HP (2016) Outlier detection methods for generalized lattices: a case study on the transition from ANOVA to REML. Theor Appl Genet 129:787–804. https://doi.org/10.1007/s00122-016-2666-6

Butler DG, Cullis BR, Gilmour AR, Gogel BG, Thompson R (2017) ASReml-R reference manual version 4. VSN International Ltd, Hemel Hempstead, HP1 1ES, UK.

Chang S, Chang T, Song Q, et al (2020) Architectural and Physiological Features to Gain High Yield in an Elite Rice Line YLY1. Rice 13:60. https://doi.org/10.1186/s12284-020-00419-y

Cullis BR, Smith A, Coombes N (2006) On the design of early generation variety trials with corrected data. J Agric Biol Environ Stat 11:381–393. https://doi.org/10.1198/108571106X154443

Cobb JN, Juma RU, Biswas PS, et al (2019) Enhancing the rate of genetic gain in public-sector plant breeding programs: lessons from the breeder’s equation. Theor Appl Genet 132:627–645. https://doi.org/10.1007/s00122-019-03317-0

Collard BCY, Gregorio GB, Thomson MJ, Islam MR, Vergara GV, Laborte AG, Nissila E, Kretzschmar T, Cobb JN (2019) Transforming rice breeding: re-designing the irrigated breeding pipeline at the international rice research institute (IRRI). Crop Breed Genet Genom 1:e190008. https://doi.org/10.20900/cbgg20190008

Damesa TM, Möhring J, Worku M, Piepho HP (2017) One step at a time: stage-wise analysis of a series of experiments. Agronomy 109:845–857. https://doi.org/10.2134/agronj2016.07.0395

Frutos E, Galindo MP, Leiva V (2014) An interactive biplot implementation in R for modeling genotype-by-environment interaction. Stoch Environ Res Risk Assess 28, 1629–1641. https://doi.org/10.1007/s00477-013-0821-z

Isik F, Holland J, Maltecca C (2017) Spatial Analysis. In: genetic data analysis for plant and animal breeding. Springer International Publishing, Cham, pp 203–226

Gerard GS, Crespo-Herrera LA, Crossa J, et al. (2020) Grain yield genetic gains and changes in physiological related traits for CIMMYT’s high rainfall wheat screening nursery tested across international environments. Field Crops Res 249:107742. https://doi.org/10.1016/j.fcr.2020.107742

Goloran JB, Johnson-Beebout SE, Morete MJ, et al (2019) Grain Zn concentrations and yield of Zn-biofortified versus Zn-efficient rice genotypes under contrasting growth conditions. Field Crops Res 234:26–32. https://doi.org/10.1016/j.fcr.2019.01.011

Hussain W, Anumalla M, Catolos M, et al (2022) Open-source analytical pipeline for robust data analysis, visualizations and sharing in crop breeding. Plant Methods 18:14. https://doi.org/10.1186/s13007-022-00845-7

Juma RU, Bartholomé J, Thathapalli Prakash P, Hussain W, Platten JD, Lopena V et al. (2021) Identification of an elite core panel as a key breeding resource to accelerate the rate of genetic improvement for irrigated rice. Rice 14, 92. https://doi.org/10.1186/s12284-021-00533-5

Khanna A, Anumalla M, Catolos M, Bartholomé J, Fritsche-Neto R, Platten JD, Pisano DJ et al. (2022a) Genetic trends estimation in IRRIs rice drought breeding program and identification of high yielding drought-tolerant Lines. Rice 15, 14. https://doi.org/10.1186/s12284-022-00559-3

Khanna A, Anumalla M, Catolos M, Bhosale S, Jarquin D Aand Hussain W (2022b) Optimizing predictions in IRRI’s rice drought breeding program by leveraging 17 years of historical data and pedigree information. Frontiers Plant Sci https://doi.org/10.3389/fpls.2022.983818

Li H, Rasheed A, Hickey LT, He Z (2018) Fast-Forwarding Genetic Gain. Trends in Plant Science 23:184–186. https://doi.org/10.1016/j.tplants.2018.01.007

Liu M, Pan T, Allakhverdiev SI, et al (2020) Crop Halophytism: An Environmentally Sustainable Solution for Global Food Security. Trends Plant Sci 25:630–634. https://doi.org/10.1016/j.tplants.2020.04.008

McLaren CG, Bruskiewich RM, Portugal AM, Cosico AB (2005) The international rice information system. a platform for meta-analysis of rice crop data. Plant Physiol 139:637–642. https://doi.org/10.1104/pp.105.063438

Merrick LF, Herr AW, Sandhu KS, et al (2022) Optimizing Plant Breeding Programs for Genomic Selection. Agronomy 12:714. https://doi.org/10.3390/agronomy12030714

Möhring J, Piepho HP (2009) Comparison of weighting in two-stage analysis of plant breeding trials. Crop Sci 49:1977–1988. https://doi.org/10.2135/cropsci2009.02.0083

Moreno-Amores J, Michel S, Miedaner T, Longin CFH, Buerstmayr H (2020) Genomic predictions for Fusarium head blight resistance in a diverse durum wheat panel: an effective incorporation of plant height and heading date as covariates. Euphytica 216:22. https://doi.org/10.1007/s10681-019-2551-x

Negrão S, Courtois B, Ahmadi N, et al (2011) Recent Updates on Salinity Stress in Rice: From Physiological to Molecular Responses. Crit Rev Plant Sci 30:329–377. https://doi.org/10.1080/07352689.2011.587725

Piepho HP, Möhring J (2007) Computing heritability and selection response from unbalanced plant breeding trials. Genetics 177:1881–1888. https://doi.org/10.1534/genetics.107.074229

Piepho HP, Möhring J, Melchinger AE, Büchse A (2008) BLUP for phenotypic selection in plant breeding and variety testing. Euphytica 161:209–228. https://doi.org/10.1007/s10681-007-9449-8

Piepho HP, Möhring J, Schulz-Streeck T, Ogutu JO (2012) A stage-wise approach for the analysis of multi-environment trials. Biom J 54:844–860. https://doi.org/10.1002/bimj.201100219

Philipp N, Weise S, Oppermann M, Börner A, Keilwagen J, Kilian B, Arend D, Zhao Y, Graner A, Reif JC, Schulthess AW (2019) Historical phenotypic data from seven decades of seed regeneration in a wheat ex situ collection. Sci Data 6:137. https://doi.org/10.1038/s41597-019-0146-y

Radanielson AM, Gaydon DS, Li T, et al (2018) Modeling salinity effect on rice growth and grain yield with ORYZA v3 and APSIM-Oryza. Eur J Agron 100:44–55. https://doi.org/10.1016/j.eja.2018.01.015

Ramstein GP, Jensen SE, Buckler ES (2019) Breaking the curse of dimensionality to identify causal variants in Breeding 4. Theor Appl Genet 132:559–567. https://doi.org/10.1007/s00122-018-3267-3

Rauf S, Shahzad M, Teixeira da Silva JA, Noorka IR (2012) Biomass partitioning and genetic analyses of salinity tolerance in sunflower (Helianthus annuus L.). J Crop Sci Biotechnol 15:205–217. https://doi.org/10.1007/s12892-011-0089-0

Rawat N, Wungrampha S, Singla-Pareek SL, et al (2022) Rewilding staple crops for the lost halophytism: Toward sustainability and profitability of agricultural production systems. Molecular Plant 15:45–64. https://doi.org/10.1016/j.molp.2021.12.003

Reynolds M, Bonnett D, Chapman SC, Furbank RT, Manès Y, Mather DE et al. (2011) Raising yield potential of wheat. I. Overview of a consortium approach and breeding strategies. J Exp Bot 62:439–452. https://doi.org/10.1093/jxb/erq311

Rutkoski JE (2019a) Estimation of Realized Rates of Genetic Gain and Indicators for Breeding Program Assessment. Crop Sci 59:981–993. https://doi.org/10.2135/cropsci2018.09.0537

Rutkoski JE (2019b) Chapter Four - A practical guide to genetic gain. In: Sparks DL (ed) Advances in Agronomy. Academic Press, pp 217–249

Sandhu N, Yadav S, Catolos M, Sta Cruz MT, Kumar A (2021) Developing climate-resilient, direct-seeded, adapted multiple-stress-tolerant rice applying genomics-assisted breeding. Front Plant Sci 12:637488. https://doi.org/10.3389/fpls.2021.637488

Singh RK, Kota S, Flowers TJ (2021) Salt tolerance in rice: seedling and reproductive stage QTL mapping come of age. Theor Appl Genet 134:3495–3533. https://doi.org/10.1007/s00122-021-03890-3

Solis CA, Yong MT, Vinarao R, et al (2020) Back to the Wild: On a Quest for Donors Toward Salinity Tolerant Rice. Front Plant Sci 11:323 https://doi.org/10.3389/fpls.2020.00323

Smajgl A, Toan T, Nhan D, Ward J, Trung NH, Tri LQ, Tri VPD, Vu PT (2015) Responding to rising sea levels in the Mekong Delta. Nature Clim Change 5:167–174. https://doi.org/10.1038/nclimate2469

Smith AB, Cullis BR, Thompson R (2005) The analysis of crop cultivar breeding and evaluation trials: an overview of current mixed model approaches. J Agric Sci 143:449–462. https://doi.org/10.1017/S0021859605005587

Smith AB, Cullis BR (2018) Plant breeding selection tools built on factor analytic mixed models for multi-environment trial data. Euphytica 214:143. https://doi.org/10.1007/s10681-018-2220-5

Yadav R, Gupta S, Gaikwad KB, et al (2021) Genetic Gain in Yield and Associated Changes in Agronomic Traits in Wheat Cultivars Developed Between 1900 and 2016 for Irrigated Ecosystems of Northwestern Plain Zone of India. Frontiers Plant Sci 12:719394. https://doi.org/10.3389/fpls.2021.719394

Yadav S, Sandhu N, Dixit S, Singh VK, Catolos M, Mazumder RR, Rahman MA, Kumar A (2021) Genomics-assisted breeding for successful development of multiple-stress-tolerant, climate-smart rice for southern and southeastern Asia. Plant Genome 14:e20074. https://doi.org/10.1002/tpg2.20074

